# Elevated levels of intracellular RNA lariats suppress the antiviral response

**DOI:** 10.1101/2024.12.07.627371

**Authors:** Chaorui Duan, Luke Buerer, Cory Bowers, Allison J. Taggart, Mara H. O’Brien, Sarah Gunasekera, Chien-Ling Lin, Jing Wang, Jonathan P. Staley, Alger M. Fredericks, Sean F. Monaghan, Anastasia Welch, Nathaniel E. Clark, Daxing Gao, Nico Marr, Shen-Ying Zhang, Jean-Laurent Casanova, William G. Fairbrother

## Abstract

Recent studies report the genetic loss of the lariat debranching enzyme (*DBR1*) activity increases susceptibility to viral infection. Here, we show that more than 25% of human introns contain large hairpin structures created by the folding of two *Alu* elements inserted in opposite orientation. In wildtype cells, this large reservoir of endogenous dsRNA is efficiently degraded. In *DBR1*-null cells, lariats accumulate in the cytosol and dsRNA becomes enriched. We demonstrate how the chronic exposure to these lariats attenuates the dsRNA sensors, reducing the response of the MDA5, RIG-I, RNase L and PKR sensing pathways. We observe evidence for both attenuation and endogenous dsRNA in anti-viral response and viral evasion. Lariats are transiently elevated during infection (e.g. HSV-1, influenza, KSHV). The HSV-1 genome expresses multiple, stable lariats that may attenuate dsRNA sensors during latency.

**HIGHLIGHTS:**

- Intronic inverted repeat *Alu* elements constitute largest source of endogenous dsRNA.
- In the absence of *DBR1*, lariats accumulate in the cytoplasm and form dsRNA.
- Chronic exposure to endogenous dsRNA in a *DBR1*-depleted environment desensitizes the dsRNA sensing pathway.
- ICP0 intron 1 of HSV-1 has a highly-structured stable lariat.

**GRAPHICAL ABSTRACT:** 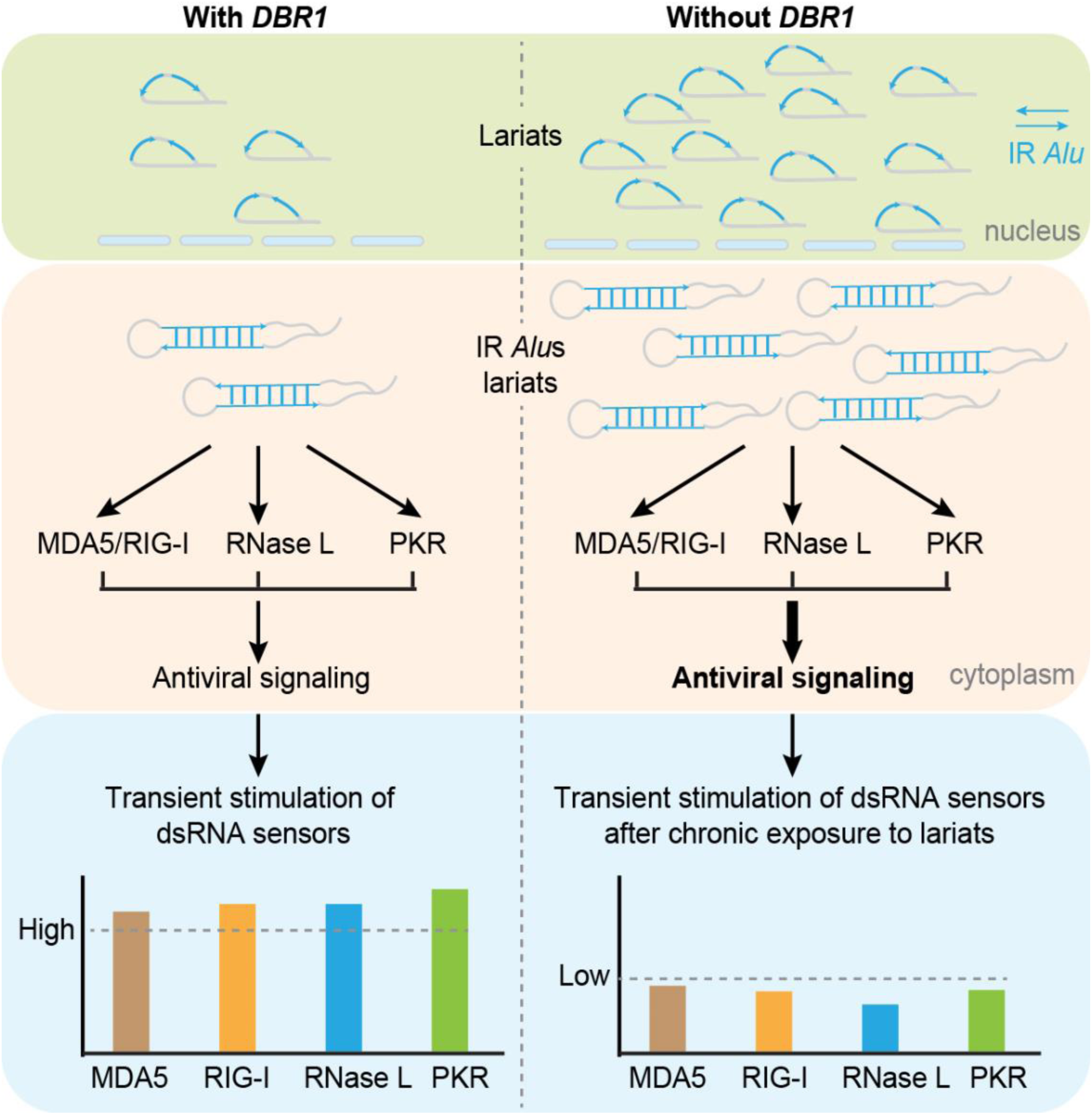

## INTRODUCTION

The excision of introns from pre-mRNA is an essential step in the production of mature mRNA transcripts. Intron removal is catalyzed by the spliceosome through two sequential nucleophilic attacks by a sugar hydroxide on the phosphate in a phosphodiester bond (Wahl et al., 2009; Wilkinson et al., 2020). First, the 5’ end of the intron forms a 2’-5’ linkage at the branchpoint to create the lariat intermediate. The second step involves a nucleophilic attack by the free hydroxyl of the upstream exon on the downstream 3’ splice site, resulting in exon ligation and the excision of the intron lariat (Padgett et al., 1984; Ruskin and Green, 1985; Ruskin et al., 1984). The vast majority of lariats are quickly linearized by the lariat debranching enzyme *DBR1*, which hydrolyzes the 2’-5’ branchpoint bond (Chapman and Boeke, 1991; Montemayor et al., 2014; Ruskin and Green, 1985). After debranching, intron lariats undergo further digestion into small RNAs or free nucleotides available for the next cycle of transcription.

Studies in mice and humans have revealed a range of pathologies caused by the loss of *DBR1* activity (Findlay et al., 2014; Zheng et al., 2015). Deficiencies in human DBR1 may lead to conditions such as cancer or impact hinder HIV replication (Galvis et al., 2014; Han et al., 2017; Xu et al., 2022; Ye et al., 2005). Bi-allelic severely hypomorphic alleles confer increased susceptibility to viral encephalitis as well as neonatal pathologies including congenital ichthyosis (Shamseldin et al., 2023; Zhang et al., 2018). The increased risk of brainstem infection can be caused by a broad spectrum of viruses, including Herpes Simplex Virus 1 (HSV-1), influenza virus, norovirus, and SARS-CoV-2 suggesting a general defect in cell intrinsic immunity (Bohmwald et al., 2021; Chan et al., 2024; Ru et al., 2024 (In press)). We previously observed elevated lariat levels upon HSV-1 infection in wildtype cells, which were enhanced by *DBR1* deficiency in the host (Zhang et al., 2018).

Fibroblasts from *DBR1*-deficient patients also displayed heightened susceptibility to both HSV-1 and Vesicular Stomatitis Virus (VSV) infections, leading to increased virus-induced cell death. These findings suggest a relationship between defects in lariat metabolism and impaired cell-intrinsic immunity. Curiously, HSV-1 and 2 encode the LAT, a highly structured lariat that resists debranching, contains dsRNA and is one of the few genes expressed during the latent phase (Farrell et al., 1991; Krummenacher et al., 1997). The specific mechanisms by which lariats modulate antiviral response has yet to be elucidated. The best-characterized antiviral RNA-sensing pathways detect cytosolic dsRNA. For example, MDA5 and RIG-I recognize viral dsRNA and trigger signaling cascades that lead to the production of interferons and pro-inflammatory cytokines, initiating an antiviral immune response (Chen and Hur, 2022; Hur, 2019). The OAS/RNase L pathway, activated by interferon signaling, degrades both viral and host RNA, promoting apoptosis and inhibiting viral replication (Cooper et al., 2014; Zhou et al., 1997). Similarly, PKR, another dsRNA sensor, phosphorylates the eukaryotic initiation factor eIF2α, resulting in global shutdown of protein synthesis and inhibition of viral replication (García et al., 2006; Hur, 2019). One of the challenges dsRNA sensors face is making a distinction between pathogenic signal and the low levels of endogenous dsRNA expressed during normal cellular function (Schlee and Hartmann, 2016).

In this study, we tested the hypothesis that stable lariat introns that accumulate in *DBR1*-deficient cells impair the responses of the cytosolic dsRNA-sensing antiviral pathways. We analyze both viral and host introns finding intronic dsRNA structure to be underrepresented in viruses and overrepresented in humans. The dominant source of secondary structure in human introns come from inverted repeat (IR) *Alu* elements. We demonstrate the induction of dsRNA sensors with acute exposure to *Alu* duplexes and attenuation of the same sensors with long term exposure to lariats. In the absence of *DBR1* activity, we visualize lariat accumulation in the cytoplasm and an increase of cytoplasmic dsRNA, which attenuates the induction of *MDA5*, *RIG-I*, and *IFNB*. There is also a loss of RNAse L activity, and lower levels of PKR and eIF2α phosphorylation in *DBR1* depleted cells which can be rescued by the add back of *DBR1*. Finally, we explore potential roles of attenuation in latent viruses. Specifically, we report the discovery of an additional structured intron in viruses (HSV-1) that is highly resistant to degradation and serves as a ligand in the PKR pathway.

## RESULTS

### Multiple viruses inhibit intron turnover in cells

Individuals with severely hypomorphic bi-allelic *DBR1* gene variants can be vulnerable to life-threatening brainstem lesions when exposed to HSV-1, Influenza virus or other viruses (Zhang et al., 2018). Previous research demonstrated that while HSV-1 infection leads to increased lariat levels in cells derived from both healthy controls and *DBR1* mutant patients, the lariat increase is significantly larger in *DBR1* mutant cells. Here we utilize influenza to infect fibroblasts derived from control and *DBR1* mutant patients to confirm that this connection between infection and decreased intron recycling can be extended across multiple viruses (Supplementary Figure S1A, 20-30% increase in lariats post infection in mutant lines, I120T *p* = 0.015 and Y17H *p* = 0.0059). Moreover, after analyzing the RNA-seq data from cells infected with Kaposi’s Sarcoma-Associated Herpesvirus (KSHV) (Najarro et al., 2024), we found that the lariats are 2.48-fold higher (Supplementary Figure S1B, *p* = 0.034) in the lytic phase compared to the latent phase. As the HSV-1, influenza and KSHV genomes are composed of intron-containing genes, it is possible that some feature of viral introns could dominantly inhibit lariat turnover. To test this hypothesis, we performed sequence analysis on the set of human introns and all the annotated introns found in all the human viral genomes.

### Viruses demonstrate adaptation towards less structured introns

We downloaded the sequences of 43 human viruses and their 126 introns from the NCBI Virus database (Hatcher et al., 2017). While GC content does not significantly differ between human and viral introns (Figure 1A), viral introns exhibit lower levels of predicted structure. On average, viral introns are shorter than human introns (9.3-fold, t-test, *p* = 7.08e-154). However, even after normalizing for length, MFE values from RNA folding predictions are lower for viral introns with the median viral intron MFE falling in the bottom 10% of the human intron MFE distribution (Figure 1B, t-test, *p* = 1.09e-09). While this result implies selection against secondary structure, we sought a measure of base pairing potential that did not rely on prediction. We calculated a maximum base pairing metric that measured the maximum proportion of positions that could be paired. Maximum base pairing potential equals 1.0 when complementary bases occur with equal frequency in the sequence (accounting for G-U wobble base pairing; see Methods). The median viral intron has 7% lower maximum base pairing potential than the median human intron, suggesting the possibility that viral sequences utilize nucleic acids in an unbalanced combination that would ensure unpaired bases and therefore less secondary structure formation (Figure 1C, t-test, *p* = 2.12e-10). These results suggest that viral introns face selective pressure towards less secondary structure, presumably due to the need to avoid triggering cytosolic dsRNA sensing mechanisms. We further interrogated the difference between the observed structure of an intron and the structure which would be expected simply based on sequence composition by generating an ensemble of 10 shuffled intron sequences for each intron and calculating the mean MFE of this shuffled ensemble (Figure 1D). Deviations between the observed and shuffled MFEs indicate introns with more or less structure than expected from their nucleotide composition. The distribution of observed and shuffled MFEs for human introns indicates that, while there is often agreement between these values, there are subsets of introns for which the observed MFE is much lower than the mean shuffled MFE (Figure 1E). In contrast, similar experiments for viral introns resulted in little difference between the observed MFEs and the expectation based on the ensemble of shuffled sequences (Figure 1F/G).

**Figure 1.**
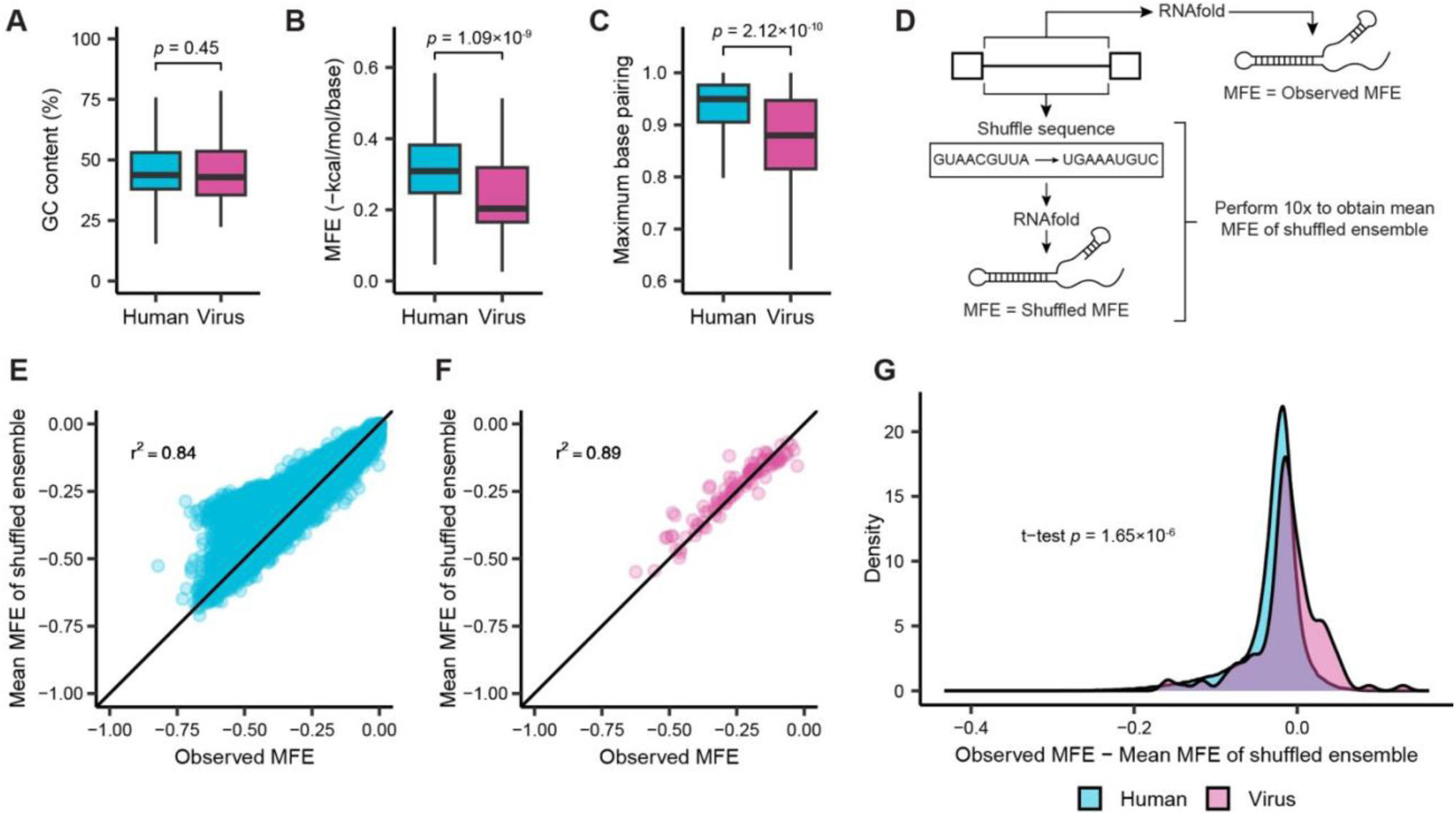
Viral introns are less structured than human introns. Human introns compared to 126 introns from human viruses according to their distributions of (A) GC content, (B) normalized predicted minimum free energy (MFE, intron length ≤ 5000 nt), and (C) potential for intron base pairing (maximum base pairing, see methods). (D) Schematic describing computational experiments comparing the observed predicted MFE of an intron to the predicted MFE of an ensemble of 10 shuffled sequences which have the same length and nucleotide sequence composition as the intron. (E) For human introns, each intron’s observed MFE is plotted against the mean MFE derived from the intron’s shuffled ensemble described in (D). (F) For human virus introns, each intron’s observed MFE is plotted against the mean MFE derived from the intron’s shuffled ensemble described in (D). (G) The distribution of the difference between observed MFE and mean MFE of shuffled ensemble for human and viral introns.

### Inverted repeat *Alu* elements contribute to human intron structure

We sought to better understand the divergence between observed and expected structure that human introns exhibited in Figure 1E. The proliferation of short interspersed nuclear elements such as *Alu* within introns could be a major contributing factor to the formation of intronic secondary structure as the presence of multiple *Alu* elements inserted within a single intron in an inverted orientation creates the potential for long stretches of base pairing due to their complementarity. To assess the relationship between *Alu* elements and intron structure, we subdivided the data into the following categories of introns containing: no *Alu*, 1 *Alu*, multiple *Alu* in the same insertion orientation, or multiple *Alu* with at least one pair in opposite insertion orientations (inverted repeat). We hypothesized that introns with inverted repeat (IR) *Alu* elements would have the largest differences between observed and shuffled MFE as the opposing orientation of *Alu* insertions in these introns provides extensive regions of near perfect complementarity. For all intron *Alu* content categories except IR *Alu*, there was a very strong correlation between the MFE of the observed intron and sequence-shuffled introns (Figure 2A). In contrast, introns with IR *Alu* had a markedly weaker correlation with many introns in this category possessing much more extensive structure than the expectation based on the ensemble of shuffled intron sequences. The ability of IR *Alu* to form long RNA hairpins within introns suggests the large fraction (28%) of introns that contain IR *Alu* would trigger cytoplasmic dsRNA surveillance pathways if they were exported to the cytoplasm. Quantifying the presence of *Alu* elements within different gene regions shows that the parts of mRNA transcripts which are exported to the cytoplasm contain fewer *Alu* than nuclear introns (Figure 2B). While the majority of introns contain *Alu*, ∼10% of 3’ untranslated regions (UTR) also contain at least one *Alu* insertion (Figure 2B). Lariat mapping of total RNA-seq samples from *DBR1* knockout and wildtype cell lines indicate lariat levels for all introns including those containing IR *Alu* increase ∼50 fold in the absence of *DBR1* activity (Figure 2C).

**Figure 2.**
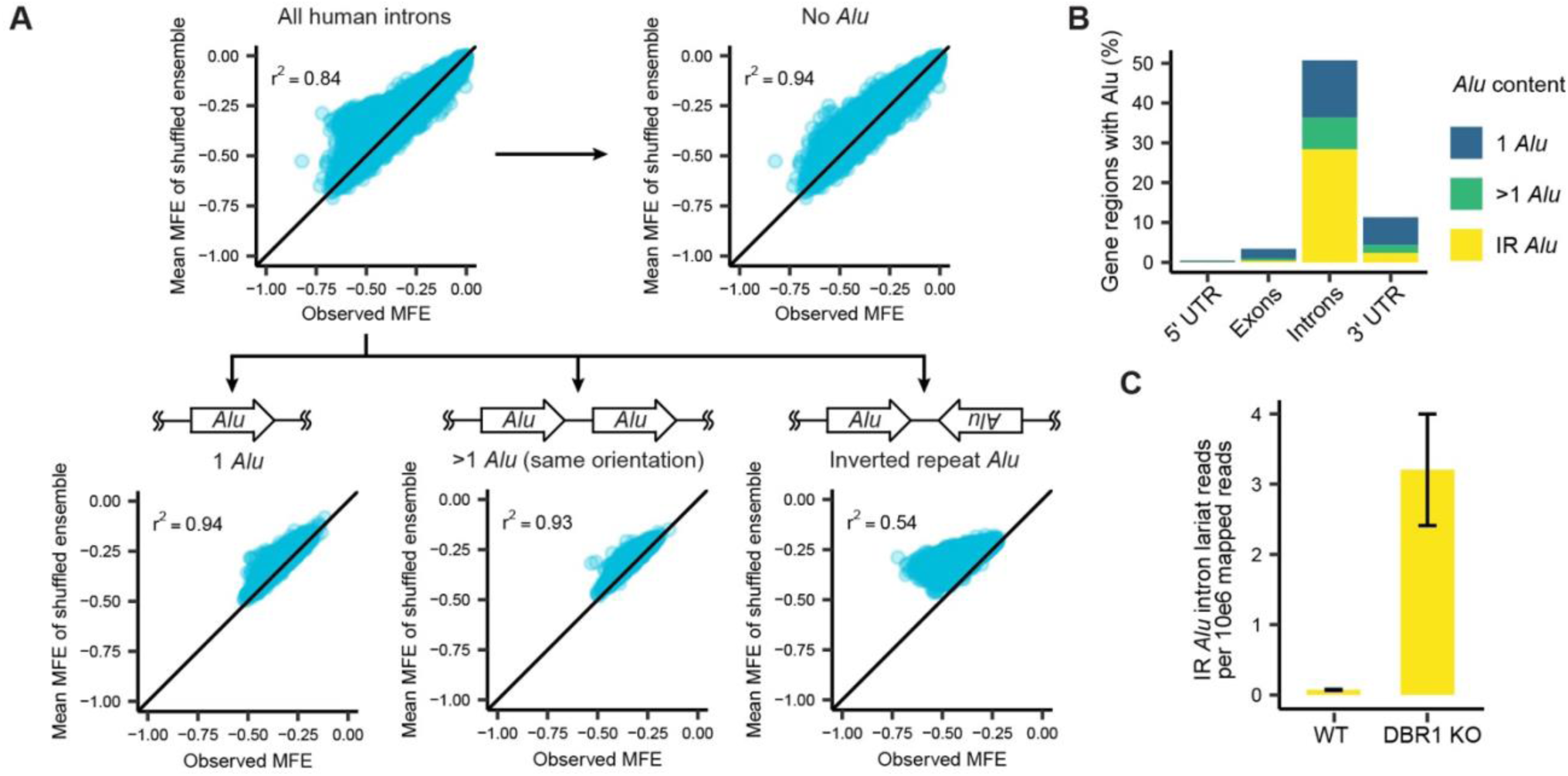
Inverted repeat *Alu* elements contribute to human intron structure. (A) The relationship between observed MFE and the mean MFE of an ensemble of shuffled intron sequences for human introns is partitioned into distinct categories based on each intron’s *Alu* content (no *Alu*, 1 *Alu*, >1 *Alu* in the same orientation, and inverted repeat (IR) *Alu*). (B) The proportion of 5’ untranslated regions (UTR), exons, introns and 3’ UTRs that contain 1 *Alu*, >1 *Alu* in the same orientation, and inverted repeat *Alu*. (C) The lariat levels of introns with IR *Alu* in wildtype and *DBR1* knockout samples.

### Lariats accumulate in the cytoplasm in the absence of *DBR1*

Given that IR *Alu* sequences can form double-stranded RNA (dsRNA) structures, the ∼50-fold increase in IR *Alu*-containing lariats suggests potential interactions with cellular dsRNA sensing mechanisms. As dsRNA sensors such as MDA5, RIG-I, PKR, TLR3, and OASes are typically situated in the cytosol(Brubaker et al., 2015; Chen and Hur, 2022; Robinson and Moehle, 2014), a key question is whether lariats that avoid debranching accumulate in the cytoplasm. To measure the distribution of lariats across different subcellular compartments, MYO19 intron 11 was chosen as an exemplar intron and its location was visualized through single molecule fluorescence in situ hybridization (FISH) performed in wildtype and *DBR1* knockout cells (intron 11, Figure 3A). Consistent with the elevated level of lariats in the absence of *DBR1*, the median FISH signal was ∼2.5-fold greater in the *DBR1* knockout cells compared to wildtype cells (Figure 3B, left). The cytoplasmic proportion of the FISH signal also increased, suggesting that lariats accumulate in the cytoplasm following a loss of *DBR1* activity (Figure 3B, right). To further investigate whether these lariats can form dsRNA in the cytoplasm, we performed immunofluorescence (IF) staining using the 9D5 antibody, which is more sensitive than the J2 antibody for dsRNA(de Faria et al., 2023). The cytoplasmic dsRNA signal was 1.4-fold higher in *DBR1* knockout cells compared to wild-type cells (Figure 3C, Mann-Whitney test, *p* = 0.0029). We also used an additional orthogonal anti-dsRNA antibody (J2) to repeat the IF assay. Co-staining confirmed that both the J2 and 9D5 antibodies detected the same cytoplasmic sites of dsRNA accumulation (Supplementary Figure S2). This result suggested that in the absence of *DBR1*, IR *Alu* lariats are exported to the cytoplasm where they form dsRNA.

**Figure 3.**
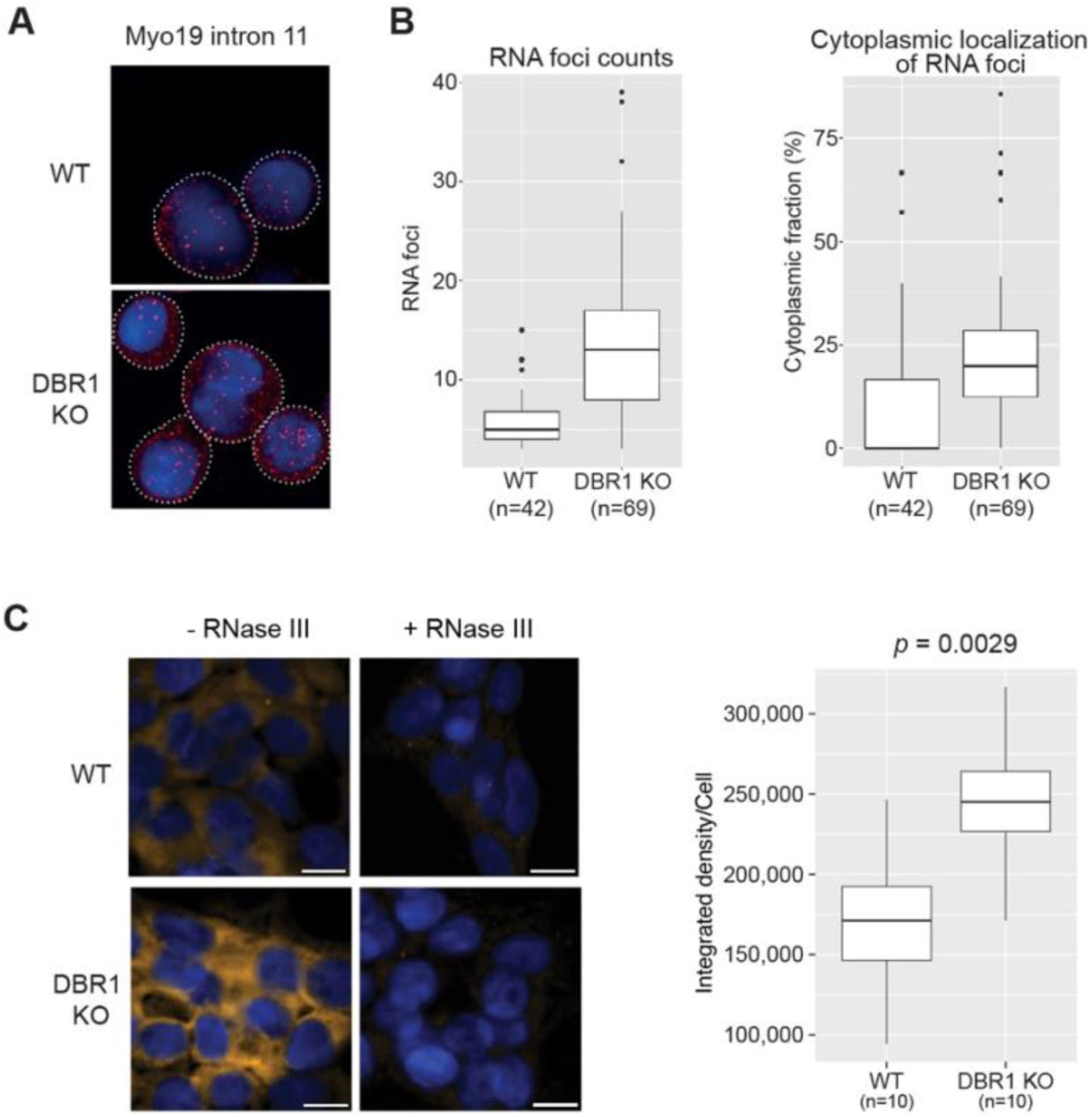
Lariats accumulate in the cytoplasm and form dsRNA in the absence of *DBR1*. (A) Imaging of FISH targeted to MYO19 intron 11 in wildtype and *DBR1* KO cells. (B) Analysis of visualized RNA foci reveals an increase in foci in *DBR1* KO samples, as well as a shift to the cytoplasm. (C) Detection of cytoplasmic dsRNA in both wild-type and *DBR1* KO cells. Cells were stained for dsRNA (9D5, yellow), with DAPI (blue) serving as a nuclear marker. Representative images of untreated and RNase III-treated cells are shown. The scale bar represents 10 µm. Quantification of cytoplasmic dsRNA signal intensity is shown on the right (*p* value from Mann-Whitney test).

### *DBR1* deficiency attenuates multiple dsRNA sensors

The accumulation of dsRNA from cytoplasmic lariats in *DBR1* knockout cells results in chronically high levels of cytoplasmic dsRNA, which is a potent stimulator of cellular dsRNA sensing mechanisms. However, the patient phenotype for *DBR1* loss of function includes a hyper-susceptibility to viral infection. In many systems, constant exposure to a ligand may result in signal attenuation via a variety of mechanisms including receptor desensitization. To test the hypothesis that chronic exposure to endogenous dsRNA in *DBR1* depleted environment desensitizes the dsRNA sensing pathway, we transfected wildtype (HEK293T) and *DBR1* knockout (C22) cells with poly I:C, a widely used mimic of the pathogenic dsRNAs present during viral infection. The real-time quantitative PCR (RT-qPCR) analysis revealed that the expression levels of the *MDA5*, *RIG-I*, and downstream interferon-beta (*IFNB*) genes in response to poly I:C were significantly reduced in C22 cells compared to HEK293T cells (Figure 4A). Specifically, *MDA5* expression was 28-33% lower, *RIG-I* was 44-57% lower, and *IFNB* was ∼75% lower in C22 cells compared to HEK293T cells. Moreover, we observed greatly reduced rRNA degradation and decreased phosphorylation levels of PKR and eIF2α in C22 cells compared to HEK293T cells (Figure 4B/C). Furthermore, upon reintroducing *DBR1* by transfecting knockout cells with a *DBR1* vector, rRNA degradation activity and PKR and eIF2α phosphorylation levels upon treatment with poly I:C were both restored (Figure 4B/D). These results suggest that the long-term elevation of lariats in *DBR1*-null cells attenuates dsRNA sensing pathways.

**Figure 4.**
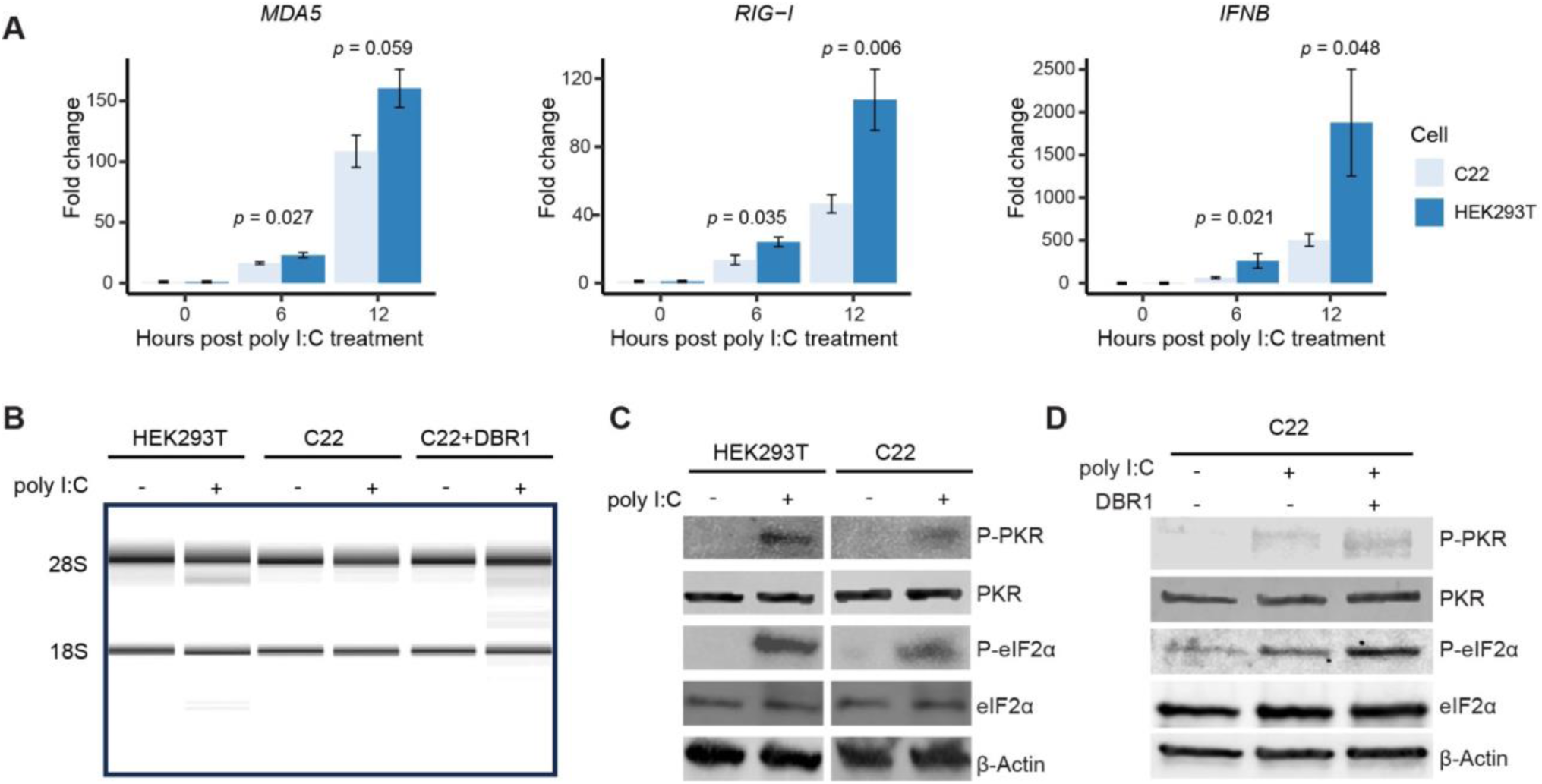
*DBR1* deficiency attenuates MDA5, RIG-I, RNase L and PKR activity. (A) mRNA expression levels of *MDA5*, *RIG-I* and *IFNB* were lower in *DBR1* knockout (KO) cells (C22) following poly I:C treatment, as determined by real-time PCR analysis. Data are shown as means from three independent experiments with standard error as error bars. *P* values were from t-test. (B) The absence of rRNA degradation was observed in *DBR1* knockout (KO) cells (C22) following 4 hours of poly I:C treatment, whereas rRNA degradation was restored upon add-back of *DBR1* (C22+DBR1) in the presence of poly I:C. (C) In *DBR1* KO cells (C22), the phosphorylation levels of PKR and eIF2α were significantly decreased compared to those in WT cells (HEK293T) following 2 hours of poly I:C treatment, as determined by western blot analysis. (D) The phosphorylation levels of PKR and eIF2α significantly increased upon add-back of *DBR1* (C22+DBR1) following a one-hour treatment with poly I:C, as determined by western blot analysis.

### *Alu* dsRNA can activate multiple dsRNA-recognizing immune sensors

IR *Alu*s are the most abundant class of cellular dsRNAs (Kim et al., 2018) and can act as immunogenic endogenous dsRNAs (Ahmad et al., 2018; Chung et al., 2018; Wu et al., 2018). To further characterize IR *Alu*s in innate immunity, we synthesized *Alu* dsRNA through in vitro transcription (*IVT*). The *IVT*-generated *Alu* dsRNA was validated using the J2 antibody. As shown by dot blot analysis, only *Alu* dsRNA is specifically detected by the J2 antibody which recognizes 40 bp of duplex RNA. RNase III treatment, which selectively degrades dsRNA, abolished the J2 signal (Figure 5A). Upon transfecting *Alu* dsRNA into wildtype cells, qPCR analysis revealed that *Alu* dsRNA can increase the expression of *MDA5* and *RIG-I* by 6 to 10-fold, and upregulate the downstream *IFNB* gene by ∼160-fold (Figure 5B). Additionally, we observed that only *Alu* dsRNA significantly induced rRNA degradation and led to increased phosphorylation of both PKR and eIF2α (Figure 5C/D).

**Figure 5.**
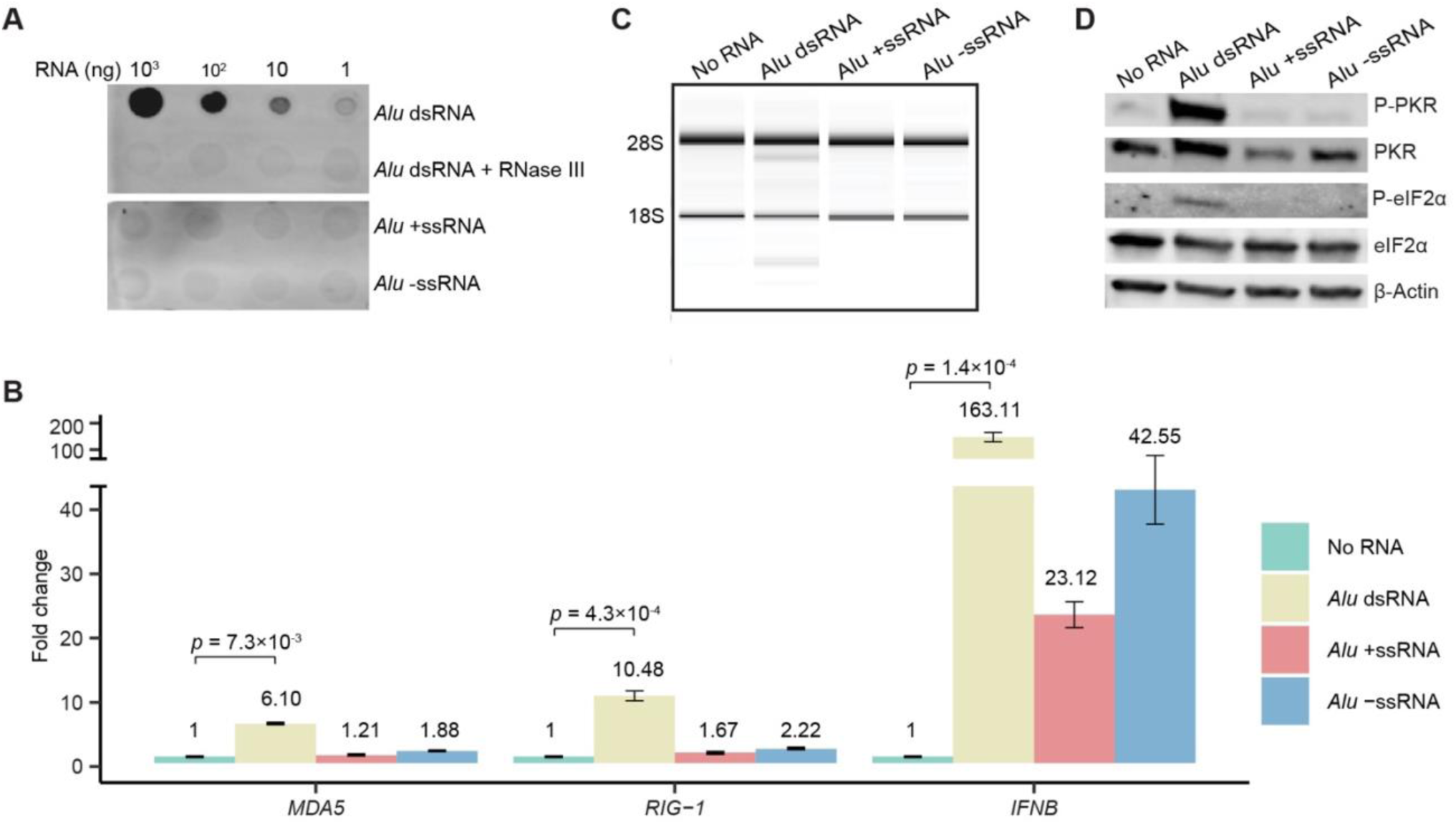
*Alu* dsRNA can activate dsRNA-recognizing immune sensors. (A) Validation of in vitro transcribed (*IVT*) *Alu* dsRNA by dot blot using the J2 antibody. (B) Wildtype cells were transfected with one or both strands of *IVT Alu* RNA for 24 hours, and qPCR analysis showed that the mRNA expression levels of *MDA5*, *RIG-I*, and *IFNB* were elevated by *Alu* dsRNA. Data are shown as means from three independent experiments with standard error as error bars. *P* values were from a two-sided t-test. (C) The Bioanalyzer analysis and (D) Western blot results showed that both rRNA degradation and the phosphorylation of PKR and eIF2α occur specifically in response to *Alu* dsRNA.

### Intron 1 of the HSV-1 ICP0 transcript is a highly structured stable lariat

The herpes simplex virus type 1 (HSV-1) has two main phases in its life cycle: lytic and latent. Studies have shown that Infected Cell Protein 0 (ICP0) plays a crucial role in initiating the lytic phase and reactivating the virus from latency (Cai et al., 1993; Halford et al., 2001; Yao and Schaffer, 1995). Given ICP0’s importance in HSV-1 infection and the fact that host RNA lariats tend to accumulate during HSV-1 infection, we became interested in what ICP0’s introns look like and how they behave during the infection process. *In silico* RNA folding prediction was performed on ICP0 introns as well as all human introns with a length ≤ 5000 nt using the RNAfold tool (Lorenz et al., 2011). ICP0 intron 1 has a predicted minimum free energy (MFE) above the 97^th^ percentile of the human intron MFE distribution (Figure 6A). The extensive regions of secondary structure in the MFE prediction likely contribute to the unusual stability of this intron in infected cells (Figure 6B). To further characterize the nature of HSV-1 intron accumulation over the course of infection, we analyzed publicly available total RNA-seq data generated from fibroblasts infected with HSV-1 for 0-8 hours (Rutkowski et al., 2015). The total RNA-seq data indicates accumulation of intronic products far exceeds that of exons after 6-8 hours post infection (Figure 6C), likely due to the stability of the ICP0 intron 1 lariat. Notably, the increased read coverage for ICP0 intron 1, compared to the previously reported stable HSV-1 latency-associated transcript (LAT) intron, implies a higher degree of stability for the ICP0 intron product. To confirm that this intronic coverage is originating from lariat RNAs, we performed lariat mapping of the total RNA-seq samples. This analysis returned many lariat reads for ICP0 intron 1 originating from lariats which utilized its 5’ splice site and a branchpoint 24 nt upstream of its 3’ splice site. The normalized lariat levels increase over the course of infection in a manner consistent with the total RNA coverage trend for this intron (Figure 6D). The highly structured and stable nature of ICP0’s lariat prompted us to investigate whether it can activate the dsRNA sensing pathway similarly to human lariats. We cloned ICP0 intron 1 along with its flanking exons into a vector and transfected it into HEK293T cell, while a vector lacking ICP0 intron 1 was transfected in parallel as a negative control. The western blot results indicated that transfection of the ICP0 intron 1 vector resulted in increased phosphorylation levels of PKR (Figure 6E).

**Figure 6.**
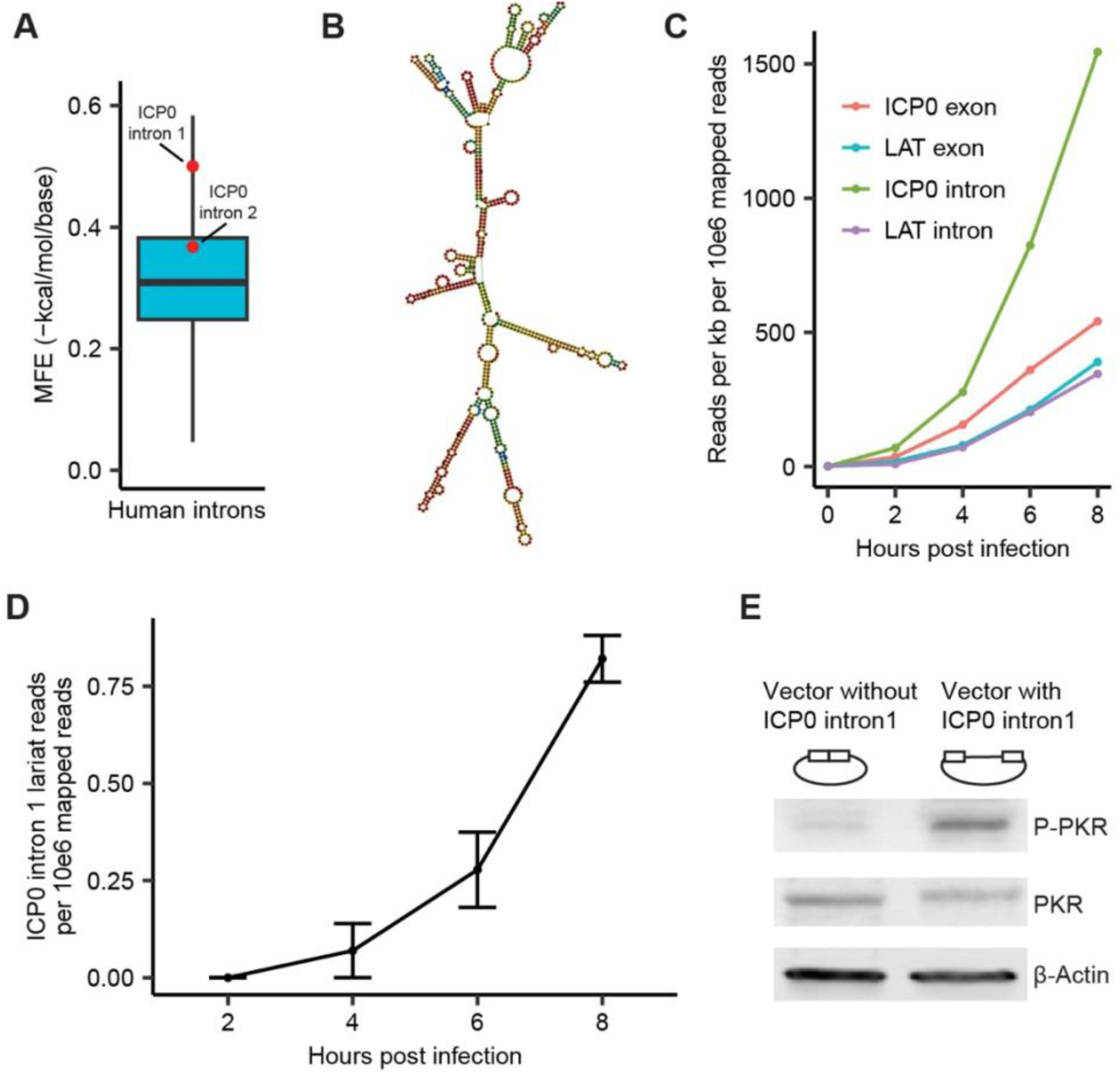
ICP0 intron 1 of HSV-1 has a highly-structured stable lariat. (A) Normalized predicted minimum free energy (MFE) for ICP0 introns 1 and 2 overlaid on the distribution of predicted MFEs for all human introns with length < 5000 nt. (B) Secondary structure of HSV-1 ICP0 stable intron 1 based on minimum free energy prediction. (C) Normalized read coverage of LAT and ICP0 intron and exon regions from total RNA-seq data of samples taken during HSV-1 infection time course experiments. (D) Normalized lariat levels for ICP0 intron 1 derived from lariat mapping of HSV-1 infection total RNA-seq data. (E) The wild-type cells were transfected with vectors containing ICP0 intron 1 and a control vector without ICP0 intron 1 for 24 hours. Western blot analysis demonstrated that the phosphorylation of PKR was specifically induced by the ICP0 intron 1 containing vector.

## DISCUSSION

We previously reported that loss of function mutations in *DBR1* impair antiviral responses to HSV-1 infections (Zhang et al., 2018). In that original report it was noted that lariat abundance increased upon HSV-1 infection regardless of *DBR1* status. Here, we extend this observation to influenza virus where cellular lariats increase upon influenza infection. Moreover, we observe that lariats increase during lytic phase after KSHV infection. These results suggest the lariat enrichment that follows viral infection could be a natural immunostimulant.

Therapeutically, this mechanism of inducing endogenous dsRNA to bolster an immune response appears to explain the efficacy of DNA methylation inhibitors in chemotherapy. These compounds derepress endogenous retroviruses which in turn induce an interferon response (Chiappinelli et al., 2015; Mehdipour et al., 2020). Our analysis suggests viruses evolve under strong selective pressure to avoid secondary structure. The observed defect in lariat recycling induced by viral infection may be an immunostimulatory signal by bringing endogenous dsRNA to the dsRNA sensors in the cytoplasm.

Initially, we explored the hypothesis that there was some property of viral introns such as increased dsRNA that acted in a dominant negative manner against the debranching machinery. However, sequence analysis of all known viral introns revealed the mark of selection against secondary structure in viral introns (Figure 1). While the GC content of human and viral introns are similar, viral introns are much less structured (Figure 1A, B).

Viral introns are encoded by an unbalanced composition of nucleotides such that at least 12% of viral positions are unpaired making it almost impossible to achieve the long stretches of double stranded required to trigger the dsRNA responses in the cytosol. In contrast, the median human host introns had only 5% of its intron unpaired (Figure 1C). Human introns sequence folded with more secondary structure than expected by chance, much of it due to hairpins that were long enough to trigger the dsRNA sensors. These ∼300 nt hairpins, found in 28% of all introns, were formed from multiple *Alu*s inserted in opposite orientations within the same intron (Figure 2). While these inverted repeats have been recognized as the major source of endogenous dsRNA, prior studies mostly focused on these elements in 3’UTR. Here we show the vast majority of transcribed IR *Alu* are intronic but are not biologically relevant because lariats are rapidly degraded.

However, in the absence of *DBR1*, there is a 50-fold increase in abundance of these endogenous dsRNA that could be recognized by the dsRNA sensors. For this recognition, lariats need to accumulate in cytoplasm (confirmed in Figure 3A, B) and contributed to increased cytoplasmic dsRNA signal (confirmed in Figure 3C and Supplementary Figure S2 by the J2 antibody that recognizes dsRNA>40bp). In the absence of *DBR1*, the lariats from these introns can escape to the cytosol and act as an endogenous source of dsRNA that interacts with cellular pathogenic dsRNA sensing. This phenomenon has also been seen with small molecular inhibition of the spliceosome and knockdown of splicing factors that presumably expose dsRNA sensors to intron duplexes comprised of inverted *Alu* repeats (Bowling et al., 2021; Zheng et al., 2024). The same intronic arrangement of *Alu* elements can reach the cytoplasm and be recognized by antibodies raised against double stranded RNA. We also demonstrate similar effects on innate immune sensors as *DBR1* knockout cells exhibited impairments in MDA5, RIG-I, RNase L and PKR responses to dsRNA (Figure 4).

Cellular accumulation of dsRNA is a common molecular signature associated with viral infections (i.e. pathogen-associated molecular pattern (PAMP), and its presence in the cell typically activates dsRNA sensors like MDA5, RIG-I and PKR, leading to an antiviral response. However, the dsRNA sensor cannot be too sensitive as there are endogenous sources of dsRNA. To add to the complexity dsRNA recognition plays a biological role during mitosis - PKR activation shuts down translation during M phase(Kim et al., 2014; Kim et al., 2018). We demonstrated that ds *Alu*s are potent ligands for recognition by multiple dsRNA sensors (Figure 5). However, in our study, we found that the long-term presence of these ligands (i.e. structured *Alu*-containing lariats) in the absence of *DBR1* attenuates dsRNA sensors like MDA5, RIG-I, RNase L and PKR. Signal attenuation after long term exposure to a ligand is a common phenomenon in receptor signaling systems(Anwar et al., 2013). While reduced sensor activity may protect the cell from the consequences of chronic endogenous dsRNA, it could also lead to increased vulnerability to viral infection. We believe this attenuation may explain the susceptibility to a broad spectrum of viruses in patients with loss of function *DBR1* alleles. Given the role of PKR activation in mitosis, the attenuated PKR we observe in the absence of *DBR1*, may result in the growth defects observed in *DBR1* KO cells (Buerer et al., 2024; Kim et al., 2014; Kim et al., 2018). While the precise mechanism of this attenuation is unclear, a recent study (Paget et al., 2023) reported that stress granules can act as cellular “shock absorbers” and protect cells from self-derived dsRNA-mediated immunopathology. Moreover, Ru et al found that the accumulation of RNA lariats in the absence of DBR1 interferes with stress granule assembly and reduces PKR activation by promoting the degradation of G3BP1 and G3BP2, which are crucial for stress granule formation (Ru et al., 2024 (In press)). Recent studies have shown that circular RNAs (circRNAs) can form short RNA duplexes that bind to PKR in an inhibitory manner (Liu et al., 2022; Liu et al., 2019). Given that lariats are a type of circular RNA, it is plausible that lariats could form a similar structure. During viral infection, these complex *Alu* lariat structures might act as a sponge, sequestering dsRNA sensors like PKR. While the precise mechanism of attenuation is not completely understood, there is some evidence that HSV-1 may utilize dsRNA to suppress dsRNA during latency. In latency, HSV-1 expresses a single gene product - a highly structured, highly stable lariat called the latency associated transcript (LAT) which is resistant to debranching (Farrell et al., 1991; Krummenacher et al., 1997). One of the speculated functions of LAT is blocking apoptosis in neurons (Perng et al., 2000). Here we discover another viral intron, ICP0 intron 1, similar to the LAT that also has a high degree of secondary structure (97^th^ percentile of normalized MFE human distribution) but has even greater stability in viral infection time course studies (Figure 6C). Transfection of a vector containing the structured intron into HEK293T activated PKR (Figure 6E).

## RESOURCE AVAILABILITY

### Lead contact

Further information and requests for reagents may be directed to and will be fulfilled by the corresponding author William G. Fairbrother.

### Materials availability

All unique materials will be available upon publication and request.

### Data and code availability

All original code has been deposited at GitHub (https://github.com/fairbrother-lab/) and is publicly available as of the date of publication.

## AUTHOR CONTRIBUTIONS

W.G.F. and J.L.C. conceived and supervised the study. C.D., M.H.O., C.L.L., J.W., A.M.F., A.W., N.E.C., N.M., D.G., and S.Y.Z. performed experiments. C.D., L.B., C.B., A.J.T., M.H.O. and S.G. performed data analysis. S.F.M provided both technical and funding support. C.D., L.B., C.B. and W.G.F. wrote the manuscript.

## ACKNOWLEDGMENTS

This work was supported by the National Institute of General Medical Sciences of the National Institutes of Health under awards R01 GM127472, R01 GM105681 and R35 GM142638.

## DECLARATION OF INTERESTS

The authors declare no competing interests. We disclose W.G.F. as the founder of Walah Scientific and serves on the scientific advisory board of Remix pharma. S.F.M. is the founder of Alcini LLC.

## STAR★METHODS

### Key resources table

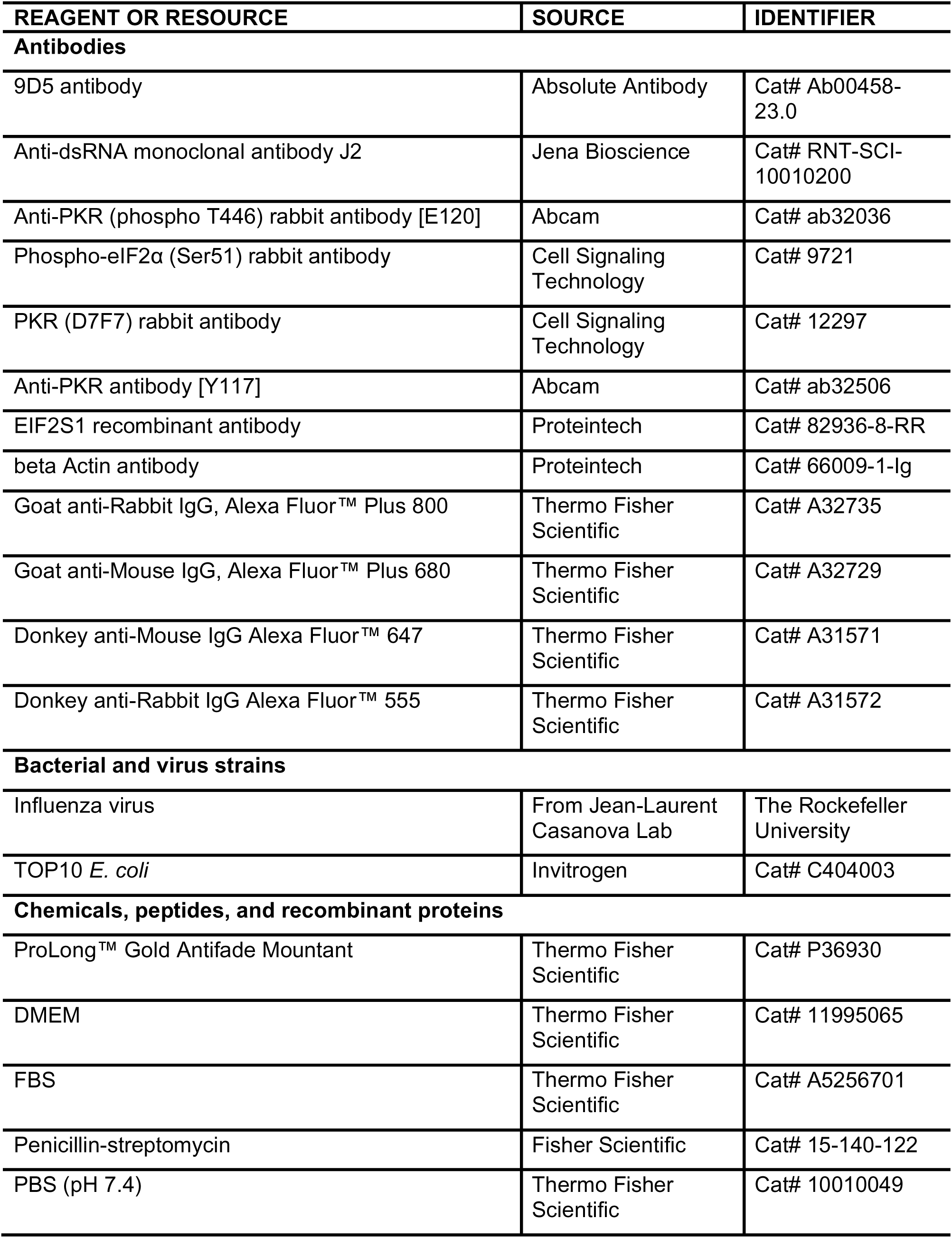
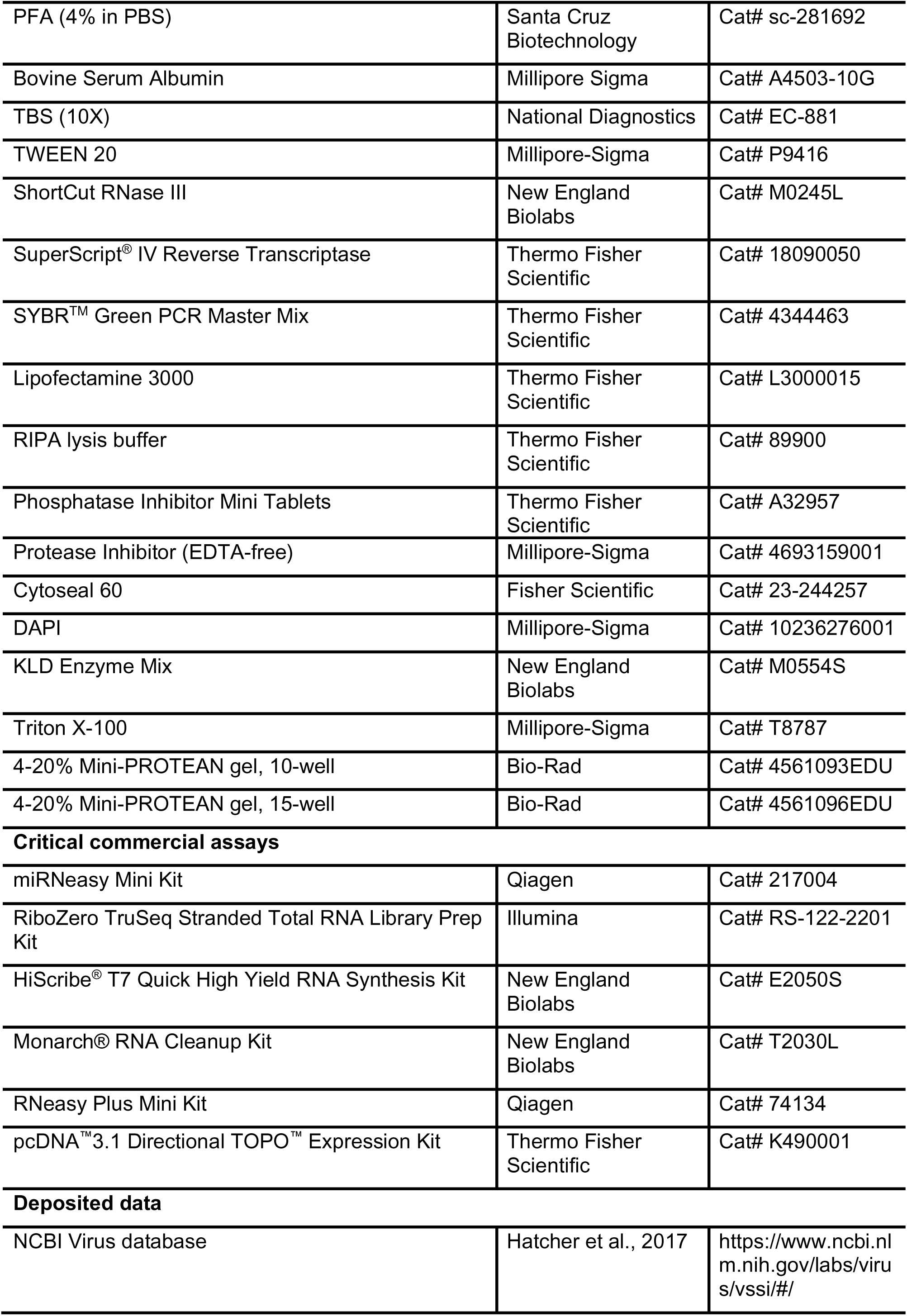
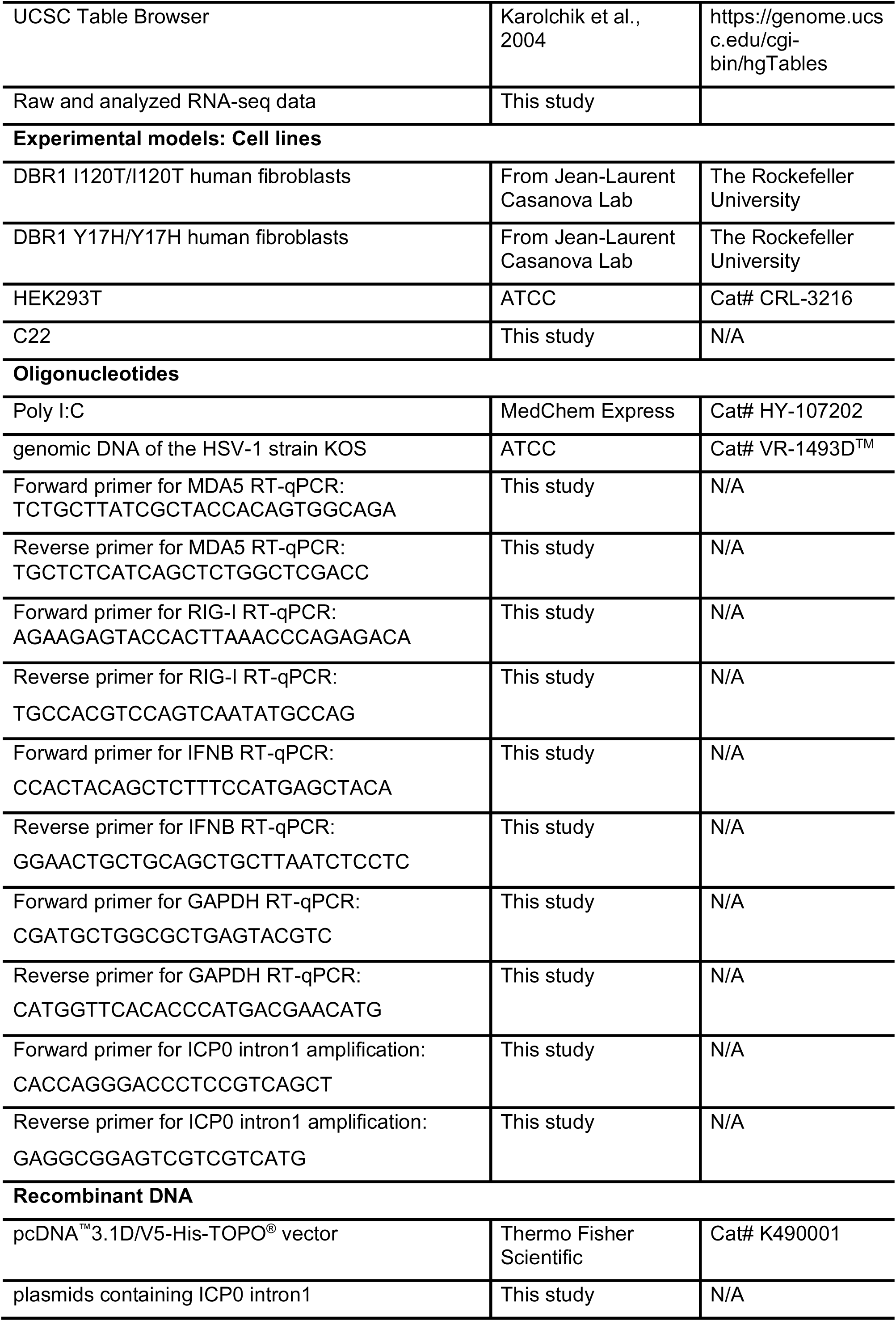
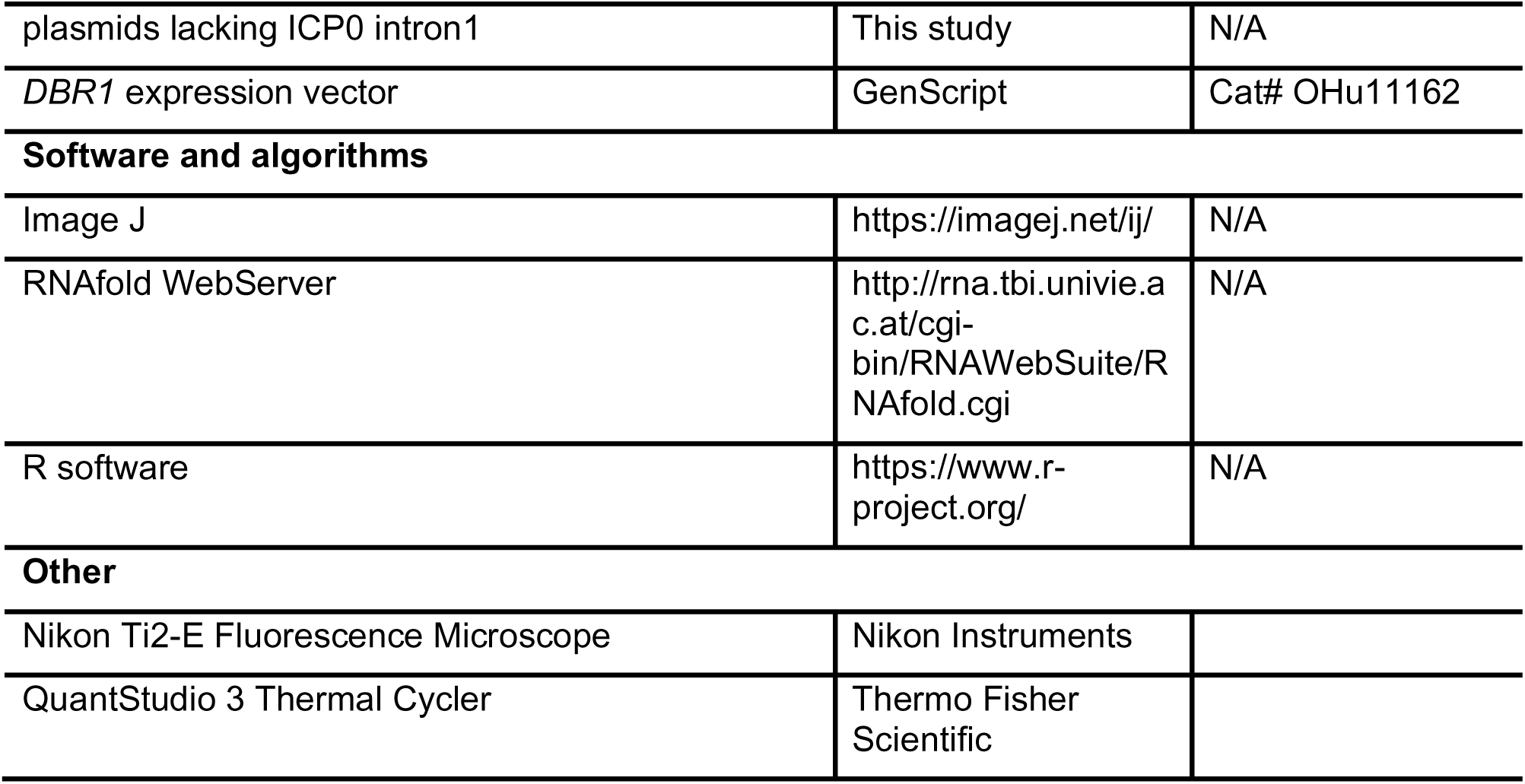

## METHOD DETAILS

### Influenza infection

*DBR1* I120T/I120T and *DBR1* Y17H/Y17H human primary fibroblasts cell lines were from a previous study (Zhang et al., 2018). For influenza infection, 4x10^5^ primary fibroblasts per well were added to 6-well plates and infected with influenza, at a multiplicity of infection (MOI) of 1, in DMEM supplemented with 2% fetal calf serum (FCS) (Everett, 1984; Gelman and Silverstein, 1985). After 2 hours, the cells were washed and incubated in 500 μL of medium. Cells were collected at 8 hours, 24 hours and frozen.

### RNA sequencing and lariat identification

Total RNA was extracted with the miRNeasy Mini Kit (QIAGEN), and treated with DNase (Ambion) to remove residual genomic DNA. RNA-seq libraries were prepared with the Illumina RiboZero TruSeq Stranded Total RNA Library Prep Kit (Illumina) and sequenced on the Illumina NextSeq platform in the 150 nt, paired-end configuration. Each library was sequenced three times.

Sequencing reads were processed using previously described lariat mapping scripts (Buerer et al., 2024). First, reads are filtered out if they contain >5% ambiguous characters. Then, reads are mapped to the genome, and aligned reads are discarded. A mapping index is then created based on the unaligned reads, and a Fasta file containing the first 20nt of each annotated intron in the transcriptome is mapped to the unaligned reads. Reads are then identified where only one 5’ splice site maps to them and the alignment has no mismatches or indels. These reads are then trimmed of the sequence from the start of the 5’ SS alignment to the end of the read, and reads where the trimmed portion is <20nt are filtered out. The remaining trimmed reads are mapped back to the genome, and the trimmed read alignments are then filtered to only consider those with <=5 mismatches, <=10% mismatch rate, and no more than one indel of <=3nt. For each trimmed read the highest scoring alignment was chosen after restricting to alignments in the same gene as the 5’ SS alignment and those with the expected inverted mapping order of the 5’ and 3’ segments. The end of this highest scoring alignment is then taken to be the branchpoint of the lariat the read is derived from.

### HSV-1 Infection and Lariats

To investigate the effects of HSV-1 infection on lariat levels, we analyzed a publicly available RNA-seq dataset from human fibroblast cells infected with HSV-1 (Rutkowski et al., 2015). This dataset was generated using HSV-1 strain 17 and a vhs-endonuclease-inactivated mutant control (both produced in baby hamster kidney (BHK) and grown in DMEM), and human foreskin fibroblasts (purchased from ECACC and cultured in DMEM). Time course data was obtained via an hourly addition of 4sU for the first 8 hours post infection (with T=0 considered after cell growth, infection, and PBS wash). This was followed by DNase treatment, RT-qPCR, library preparation and sequencing (Illumina), and RNA sequencing data mapping (ContextMap). Stranded RNA-seq reads (2 x 101nt) allowed for comparison of reads derived from the HSV-1 LAT versus reads derived from the HSV-1 ICP0 intron. Lariat mapping was performed with the algorithm described above.

### Viral Intron Analysis

Sequence data for viral introns were obtained via the NCBI Virus database (’RefSeq’ > ‘Homo sapiens (human), taxid:9606’) (Hatcher et al., 2017). The locations and sequences were extracted from the GenBank data. This dataset was filtered to remove duplicates, resulting in a final dataset of 127 viral introns. Sequence data for 210,290 human introns from the CCDS annotation were obtained via the UCSC Table Browser tool (’Feb. 2009 (GRCh37/hg19’ > ‘Genes and Gene Predictions’ > ‘CCDS’ > ‘ccdsGene’) (Karolchik et al., 2004).

To limit the computation required for in silico folding prediction, only the 159,121 human introns with length ≤ 5000nt were processed with RNAfold. These introns as well as all viral introns were processed with RNAfold using default options (Lorenz et al., 2011).

Maximum Base Pairing (MBP) can be calculated as follows:

A, T, C, G = proportions of each nucleotide in intron sequence

MBP = 1 – ( max(A-T, 0) + max(C-G, 0) + | max(T-A, 0) - max(G-C, 0) |)

ex: sequence = ATCGCGAT

A = 0.25 T = 0.25 C = 0.25 G = 0.25

MBP = 1 – ( max(0.25-0.25, 0) + max(0.25-0.25, 0) + | max(0.25-0.25, 0) – max(0.25-0.25, 0) |)

MBP = 1 – (0 + 0 + | 0 - 0 |) = 1

ex: sequence = AAAACCCC

A = 0.5 T = 0 C = 0.5G = 0

MBP = 1 – ( max(0.5-0, 0) + max(0.5-0, 0) + | max(0-0.5, 0) – max(0-0.5, 0) |)

MBP = 1 – (0.5 + 0.5 + | 0 - 0 |) = 0

ex: sequence = AAAATTCC

A = 0.5 T = 0.25 C = 0.25 G = 0

MBP = 1 – ( max(0.5-0.25, 0) + max(0.25-0, 0) + | max(0.25-0.5, 0) – max(0-0.25, 0) |)

MBP = 1 – (0.25 + 0.25 + | 0 - 0 |) = 0.5

### Fluorescence in-situ hybridization

To visualize lariat RNA species from intron 11 of the MYO19 gene, Quasar570^®^-labeled smRNA-FISH probes were ordered from Stellaris RNA-FISH (LGC Biosearch technologies). smRNA-FISH was performed according to the manufacturer’s instructions. Briefly, 1 x 10^6^ cells were fixed in 3.7% (vol./vol.) formaldehyde and then permeabilized in 70% ethanol at 4°C. For in situ hybridization, permeabilized cells were incubated with 1.25 µM probes in the hybridization buffer (10% formamide, 10% dextran sulfate, 1 mg/mL E. coli RNA, and 0.2 mg/mL BSA in 2X SSC) at 37°C overnight. Afterward, cells were washed sequentially with wash buffer A (10% formamide in 2X SSC), wash buffer B (0.1% triton-x-100 in 2X SSC), wash buffer C (1X SSC with 1 µg/mL DAPI), and 1X PBS. Finally, these cells were plated onto the coverslip and mounted in ProLong™ Gold Antifade Mountant (Thermo Fisher Scientific). The mounted cells were imaged using an Olympus IX81 inverted widefield microscope equipped with Hamamatsu Orca Flash 4.0 camera with 4 megapixels and a 100x 1.45NA oil objective lens. Single RNA molecule counting was done using Image J, and statistical analysis was conducted in R.

### Immunofluorescence

Glass coverslips were sterilized with 70% ethanol and dried under ultraviolet light. Cells were seeded in 6-well plates (300,000 cells per well) and grown on coverslips for 48 hours in DMEM containing 10% fetal bovine serum (FBS), and 100 U/ml penicillin-streptomycin (P/S). Media was removed, cells were rinsed in pre-warmed PBS, fixed for 10 minutes using 4% PFA/PBS, washed in PBS, permeabilized for 10 minutes using 0.1% TritonX-100/ PBS, washed in PBS-T, washed in blocking buffer (1% BSA/TBS-T) for 10 minutes, and treated for 3 hours at 37°C with either reaction mixture alone, or with RNase III (New England Biolabs) using 50 U per well, supplied buffers, and supplemented with 3 mM magnesium chloride. Cells were then washed in TBS-T and blocked (1% BSA/TBS-T) for one hour. Primary antibody incubation was carried out overnight at 4°C using 9D5 (Absolute Antibody, Ab00458-23.0) diluted to 1 µg/mL in blocking buffer. Cells were washed in TBS-T and treated at room temperature for 45 minutes with secondary antibody Donkey anti-Rabbit IgG, Alexa Fluor™ 555 (Thermo Fisher Scientific, A31572) and Donkey anti-Mouse IgG, Alexa Fluor™ 647 (Thermo Fisher Scientific, A31571) and 450 nM DAPI (Millipore-Sigma) in blocking buffer. Cells were washed in TBS-T and mounted to slides using Cytoseal 60 (Fisher Scientific) mounting medium.

### Microscopy and image analysis

Cells were imaged at 60x magnification using a Nikon Ti2-E Fluorescence Microscope (Nikon Instruments, Inc). DAPI and TRITC filters were used to measure nuclear signal and dsRNA signal, respectively. Signal intensities were quantified using Image J software. Following background subtraction, cells were counted using auto-thresholding, watershed segmentation, and particle analysis. Larger-sized particles were visually inspected to determine the actual cell number. Cytoplasmic integrated density was determined by subtracting TRITC staining occurring in nuclear regions from total signal, and then dividing by the total number of cells.

### *Alu* dsRNA synthesis, validation and transfection

*Alu* dsRNA was synthesized in vitro using the HiScribe^®^ T7 Quick High Yield RNA Synthesis Kit (New England Biolabs). To create the DNA template for *Alu* dsRNA, the T7 promoter sequence (5’-TAATACGACTCACTATAGGG-3’) was added to the 5’ end of both primers. For the single-stranded sense and antisense *Alu* templates, the T7 promoter sequence was added to the 5’ end of either the forward or reverse primer. The synthesized RNAs were treated with DNase to remove the DNA template, followed by purification using the Monarch® RNA Cleanup Kit (New England Biolabs).

The RNAs were then serially diluted and dotted onto a nylon membrane. In parallel, the *Alu* dsRNA was treated with RNase III (New England Biolabs) according to the manufacturer’s protocol. The RNAs were crosslinked to the membrane using UV light at 125 mJ/cm² at 254 nm. The blot was probed with the Anti-dsRNA monoclonal antibody J2 (Jena Bioscience, RNT-SCI-10010200).

5 µg of each single-stranded sense *Alu* RNA (*Alu* +ssRNA) and antisense *Alu* RNA (*Alu*-ssRNA), as well as *Alu* dsRNA, were transfected into HEK293T cells in triplicate. After 24 hours, RNAs and proteins were extracted for subsequent analyses, including real-time quantitative PCR, Bioanalyzer profiling, and western blotting.

### Poly I:C treatment

Cells were seeded into a 6-well plate and cultured in DMEM containing 10% FBS and 100 U/ml P/S. Before poly I:C transfection, the cell culture medium was replaced with serum-free medium. Poly I:C (MedChem Express, HY-107202) at a concentration of 10 μg/ml was transfected using Lipofectamine 3000 (Invitrogen) in serum-free medium and incubated at 37°C. After 4 hours, cells were collected, and RNA was extracted using the RNeasy Plus Mini Kit (Qiagen), followed by Bioanalyzer analysis (Agilent 2100). 6 hours after transfection, the medium in each well was replaced with a fresh serum-containing medium. Cells were collected 6 and 12 hours after transfection for real-time quantitative PCR analysis.

### Construction and Transfection of the Vector Containing ICP0 Intron 1

ICP0 intron 1 along with its flanking exons were cloned from genomic DNA of the HSV-1 strain KOS (ATCC^®^ VR-1493D^™^) via PCR using the forward primer (5’-3’) CACCAGGGACCCTCCGTCAGCT and reverse primer (5’-3’) GAGGCGGAGTCGTCGTCATG. The amplified ICP0 intron 1 with its flanking exons was then inserted into the pcDNA^™^3.1D/V5-His-TOPO^®^ vector using the pcDNA^™^3.1 Directional TOPO^™^ Expression Kit (Thermo Fisher Scientific). To remove ICP0 intron 1, inverted PCR was performed using the forward primer (5’-3’) CCCGCCCCGGATGTCTGG and the reverse primer (5’-3’) CTCGCGCTGGGGGCGG, followed by a KLD reaction (New England Biolabs) according to the manufacturer’s protocol. The final constructs were confirmed by whole plasmid sequencing (Eurofins). Subsequently, 5 µg of plasmids containing ICP0 intron1 and plasmids lacking ICP0 intron1 were transfected into HEK293T cells in triplicate. RNA and protein were extracted 24 hours post-transfection for analysis using real-time PCR, Bioanalyzer, and western blot.

### Real-time quantitative PCR

The total RNA was isolated from cells using the RNeasy Plus Mini Kit (Qiagen). cDNA synthesis was carried out using SuperScript^®^ IV Reverse Transcriptase (Invitrogen). RT-qPCR reactions were performed on a QuantStudio 3 Thermal Cycler with SYBR^TM^ Green PCR Master Mix (Thermo Scientific). The mRNA expression levels were calculated using the 2-ΔΔCt method (Livak and Schmittgen, 2001) and normalized to the reference gene *GAPDH*.

### Western Blot

Cells were seeded into a 6-well plate with serum-free medium and then incubated overnight. Poly I:C was transfected at a concentration of 10 μg/ml using Lipofectamine 3000 (Invitrogen) in serum-free medium and incubated at 37°C for 2 hours. For *DBR1* add-back, 5 μg of *DBR1* expression vector (GenScript, OHu11162) was transfected into cells using Lipofectamine 3000 (Invitrogen) 24 hours prior to the poly I:C treatment. Cell lysates were prepared using radioimmunoprecipitation assay (RIPA) lysis buffer containing protease and phosphatase inhibitors. Protein samples were separated on 4-20% Mini-PROTEAN gel (Bio-Rad) and transferred to a polyvinylidene difluoride (PVDF) membrane. The blot was then probed with Anti-PKR (phospho T446) rabbit antibody [E120] (Abcam, ab32036), Phospho-eIF2α (Ser51) rabbit antibody (Cell Signaling Technology, 9721), PKR (D7F7) rabbit antibody (Cell Signaling Technology, 12297)/Anti-PKR antibody [Y117] (Abcam, ab32506), EIF2S1 recombinant antibody (Proteintech, 82936-8-RR) and beta Actin antibody (Proteintech, 66009-1-Ig). Goat anti-Rabbit IgG (H+L) Highly Cross-Adsorbed Secondary Antibody, Alexa Fluor™ Plus 800 (Thermo Fisher Scientific, A32735) and Goat anti-Mouse IgG (H+L) Highly Cross-Adsorbed Secondary Antibody, Alexa Fluor™ Plus 680 (Thermo Fisher Scientific, A32729) were used according to the manufacturer’s instructions. The Bio-Rad imager was used to capture the images.

**Supplementary Figure S1.**
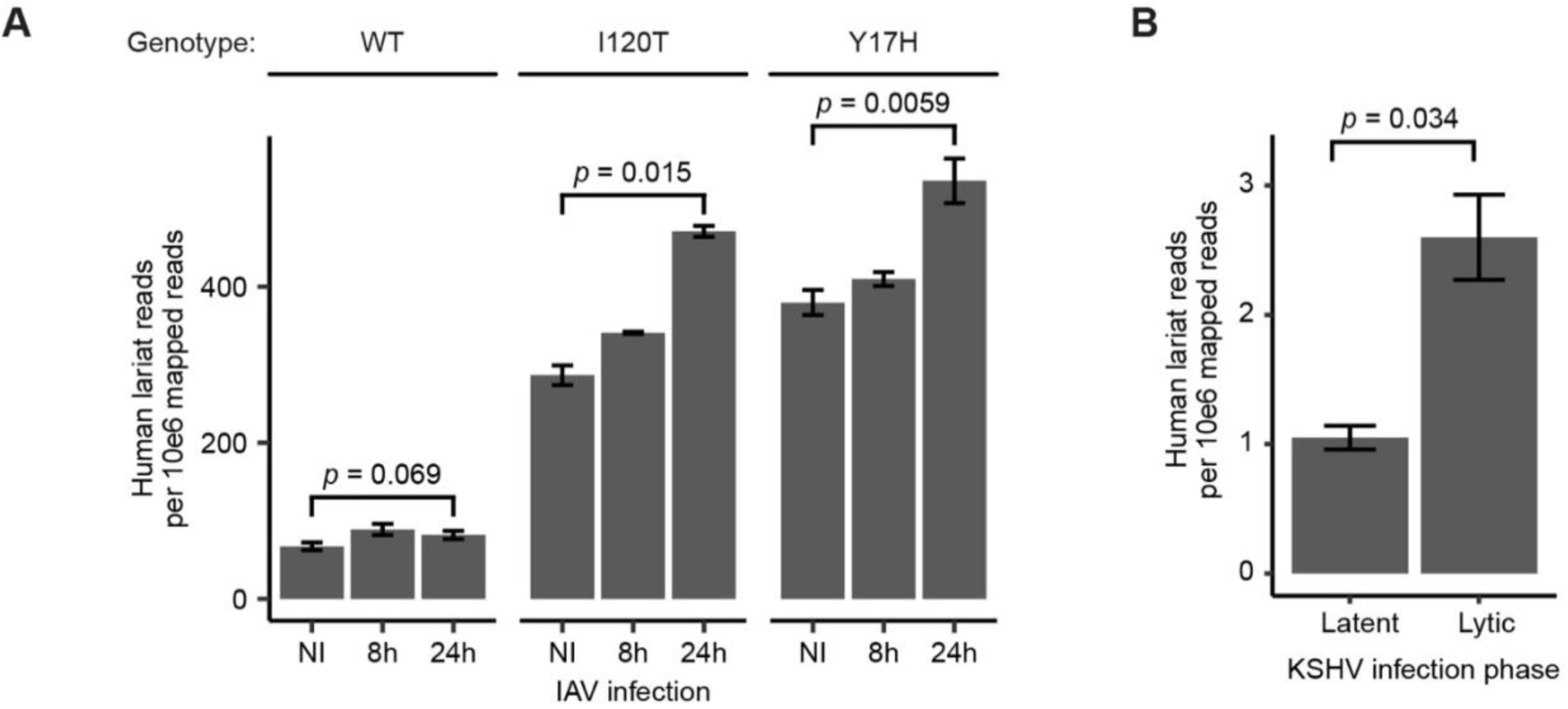
Cellular lariats increase upon virus infection. **(A) Cellular lariats increase in *DBR1* mutant cells upon Influenza infection.** Human lariat levels in RNA-seq data from wildtype (WT) and *DBR1* mutant (I120T and Y17H) patient cell lines that were not infected (NI), infected with influenza virus (IAV) for 8 hours (8h), and infected with IAV for 24 hours (24h) (*p* value from two-sided t-test). **(B) Cellular lariats increase during lytic phase upon KSHV infection.** Human lariat levels during the latent and lytic phases in RNA-seq data from cells infected with KSHV for 48 hours.

**Supplementary Figure S2.**
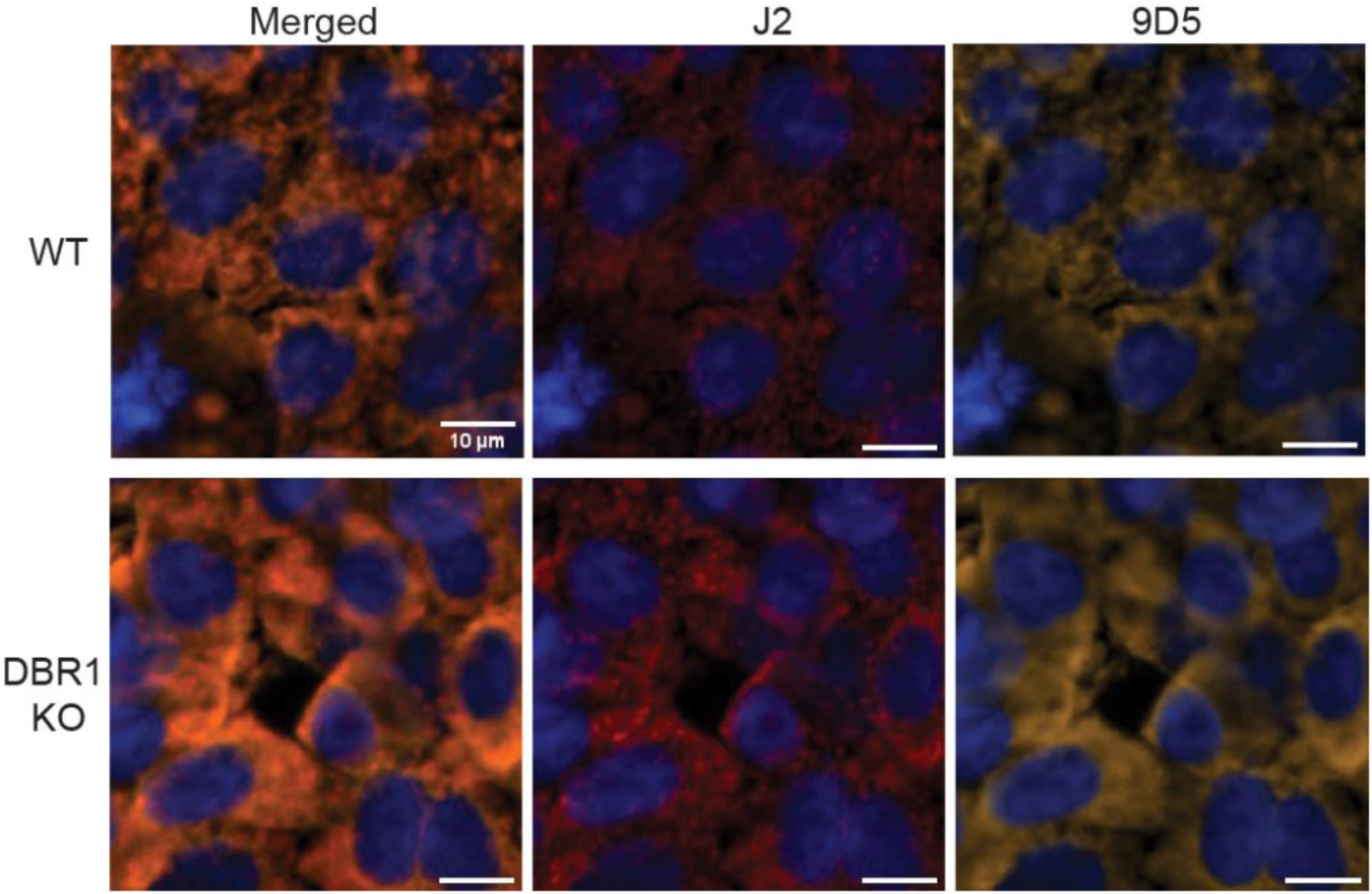
J2 (red) and 9D5 (yellow) dsRNA antibodies stain the same foci within wildtype and *DBR1* knockout cells.

